# BCL-XL is essential for the protection from secondary anemia caused by radiation-induced fatal kidney damage

**DOI:** 10.1101/2020.04.28.055665

**Authors:** Kerstin Brinkmann, Paul Waring, Stefan Glaser, Verena Wimmer, Duong Nhu, Lachlan Whitehead, Alex RD Delbridge, Guillaume Lessene, Marco J Herold, Gemma L Kelly, Stephanie Grabow, Andreas Strasser

## Abstract

Studies of gene-targeted mice identified the roles of the different pro-survival BCL-2 proteins during embryogenesis, but less is known about the roles of these proteins in adults, including in the response to cytotoxic stresses, such as treatment with anti-cancer agents. We investigated the role of BCL-XL in adult mice using a strategy where prior bone marrow transplantation allowed for loss of BCL-XL exclusively in non-hematopoietic tissues to prevent anemia caused by BCL-XL-deficiency in erythroid cells. Unexpectedly, the combination of total-body γ-irradiation (TBI) and genetic loss of *Bcl*-*x* caused secondary anemia resulting from chronic renal failure due to apoptosis of renal tubular epithelium with secondary obstructive nephropathy. These findings identify a critical protective role of BCL-XL in the adult kidney and inform on the use of BCL-XL inhibitors in combinations with DNA damage-inducing drugs for cancer therapy.

**Summary:** The inducible loss of BCL-XL in all cells of adult mice causes primary anemia due to apoptosis of erythroid and megakaryocytic cell populations. In contrast γ-radiation plus loss of BCL-XL in all cells except hematopoietic cells causes secondary anemia resulting from kidney damage.

## Introduction

Evasion of apoptotic cell death is a hallmark of cancer (Hanahan and Weinberg, 2000; Hanahan and Weinberg, 2011) and direct activation of the cell death machinery by BH3-mimetic drugs represents an attractive therapeutic strategy (Merino et al., 2018). Apoptosis is controlled by pro- and anti-apoptotic members of the BCL-2 protein family (Tait and Green, 2010), with the transcriptional or post-transcriptional activation of the pro-apoptotic BH3-only members of the BCL-2 family (e.g. BIM, PUMA) being its initiator. They bind with high affinity to the anti-apoptotic BCL-2 family members (e.g. BCL-2, BCL-XL), thereby unleashing the pro-apoptotic effector proteins BAX and BAK to cause mitochondrial outer membrane permeabilization (MOMP). MOMP causes the release of activators of the caspase cascade that causes cell demolition (Riedl and Salvesen, 2007; Tait and Green, 2010). The extent of MOMP and apoptosis are considered key determinants of therapeutic success of anti-cancer therapeutics (Del Gaizo Moore and Letai, 2013). BH3-only proteins, particularly PUMA (Villunger et al., 2003) and BIM (Bouillet et al., 1999a), are critical for the killing of malignant (and non-transformed) cells by diverse anti-cancer agents. As a consequence of genetic or epigenetic changes, many cancers show a reduction in pro-apoptotic BH3-only proteins or over-expression of pro-survival BCL-2 proteins (Beroukhim et al., 2010; Uhlen et al., 2005; Uhlen et al., 2017). Certain cancer cells can be killed by the loss of a single pro-survival BCL-2 family member (Glaser et al., 2012; Kelly et al., 2014), whereas others require loss/inhibition of two or more of these proteins (Campbell and Tait, 2018). Several small-molecule inhibitors of pro-survival BCL-2 proteins (BH3-mimetic drugs) have been developed (Adams and Cory, 2018). Even though BH3-mimetics can potently kill diverse cancer cells *in vitro* and *in vivo*, the safe clinical use of some of these drugs remains challenging due to the obligate roles of pro-survival BCL-2 proteins for the survival of normal cells in healthy tissues (Adams and Cory, 2018).

Gene deletion studies in mice have helped identify the critical roles of the different pro-survival BCL-2 family members. Loss of A1/BFL-1 causes minor reductions in certain hematopoietic cell subsets (Schenk et al., 2017; Tuzlak et al., 2017) and loss of BCL-W causes male sterility (Print et al., 1998). BCL-2-deficient mice die soon after weaning due to polycystic kidney disease and also present with reductions in mature B as well as T lymphocytes and melanocytes (causing premature greying) (Veis et al., 1993). These abnormalities could be prevented by concomitant loss of pro-apoptotic BIM (Bouillet et al., 2001). Constitutive loss of MCL-1 causes embryonic lethality prior to implantation (E3.5) (Rinkenberger et al., 2000) and tissue-restricted deletion revealed that many cell types, including cardiomyocytes (Thomas et al., 2013; Wang et al., 2013), neurons (Arbour et al., 2008) and several hematopoietic cell subsets (Opferman et al., 2005) require MCL-1 for survival. Loss of *Bcl*-*x* causes embryonic lethality at ~E13.5 with aberrant death of neurons and immature hematopoietic cells (Motoyama et al., 1995). Conditional deletion of *Bcl*-*x* in erythroid cells causes fatal anemia (Wagner et al., 2000) and loss of just one allele of *Bcl*-*x* impairs male fertility (Rucker et al., 2000) and reduces platelet numbers (Mason et al., 2007). BH3-mimetic drugs that selectively target BCL-XL (A1331852) (Lessene et al., 2013) or BCL-2 (ABT-199) *(Souers et al., 2013)* or compounds that inhibit BCL-XL, BCL-2 and BCL-W (ABT-737, ABT-263) (Oltersdorf et al., 2005; Tse et al., 2008) have been developed. Even though inhibitors of BCL-XL can efficiently kill diverse cancer cells, by themselves or in combination with additional anti-cancer agents (Cragg et al., 2008; Cragg et al., 2007; Oltersdorf et al., 2005; Tse et al., 2008), these compounds are progressing slowly in the clinic. This is in part due to the fact that the essential roles of BCL-XL in the adult (beyond its role in hematopoietic cells) are not clearly understood. Here we investigated the impact of inducible loss of BCL-XL in adult mice. This revealed that BCL-XL loss is tolerated throughout the body as long as the hematopoietic cells remain BCL-XL-sufficient. However, mice that had been γ-irradiated (for hematopoietic transplantation) and then subjected to loss of BCL-XL developed secondary anemia due to chronic renal failure characterized by renal tubular epithelial cell apoptosis with secondary obstructive nephropathy. This morbidity could be markedly delayed, although not abrogated, by the concomitant loss of pro-apoptotic PUMA or BIM. These findings predict potential toxicities and their underlying mechanisms resulting from combinations of BCL-XL inhibitors with DNA damage-inducing anti-cancer therapeutics.

## Results

### Impact of induced deletion of BCL-XL in adult mice

*Bcl-x*^−/−^ embryos die ~E13.5 due to loss of neuronal and erythroid cells (Motoyama et al., 1995). Mice with hypo-morphic mutations in the *Bcl*-*x* gene are viable but develop severe thrombocytopenia and anemia *(Mason et al., 2007)*. We wanted to explore the consequences of inducible loss of BCL-XL in adult mice. For this, we crossed *Bcl-x*^*fl*/*fl*^ mice with *RosaCreERT2* mice that express a CreERT2 fusion protein in all tissues. CreERT2 is normally kept inactive in the cytoplasm by binding to HSP90 but can be activated by tamoxifen (Vooijs et al., 2001). Even though BCL-XL is essential for neuronal cell survival (Motoyama et al., 1995), we did not expect any neuronal abnormalities upon tamoxifen-treatment in *Bcl-x*^*fl*/*fl*^;*RosaCreERT2*^+/*Ki*^ mice since tamoxifen administered by oral gavage did not efficiently drive CreERT2-mediated deletion of the *Bcl*-*x*-floxed genes in the brain, as shown by Southern blotting (Supplementary Figure S1A). Almost complete deletion of the *Bcl*-*x* gene was achieved in most other tissues, including the spleen, kidney, liver, pancreas, intestine and lung, whereas deletion in the testis was ~70%. As reported (Motoyama et al., 1995; Wagner et al., 2000), we found that induced loss of BCL-XL in adult mice caused fatal thrombocytopenia and anemia with a median survival of ~25 days (Figure 1A). Accordingly, the BCL-XL-deleted mice presented with significant decreases in platelets, red blood cells (RBCs), hemoglobin content (HGB) and hematocrit (HCT) (Figures 1C-F), accompanied by massively enlarged spleens due to compensatory erythropoiesis (Figure 1G-I).

**Figure 1:**
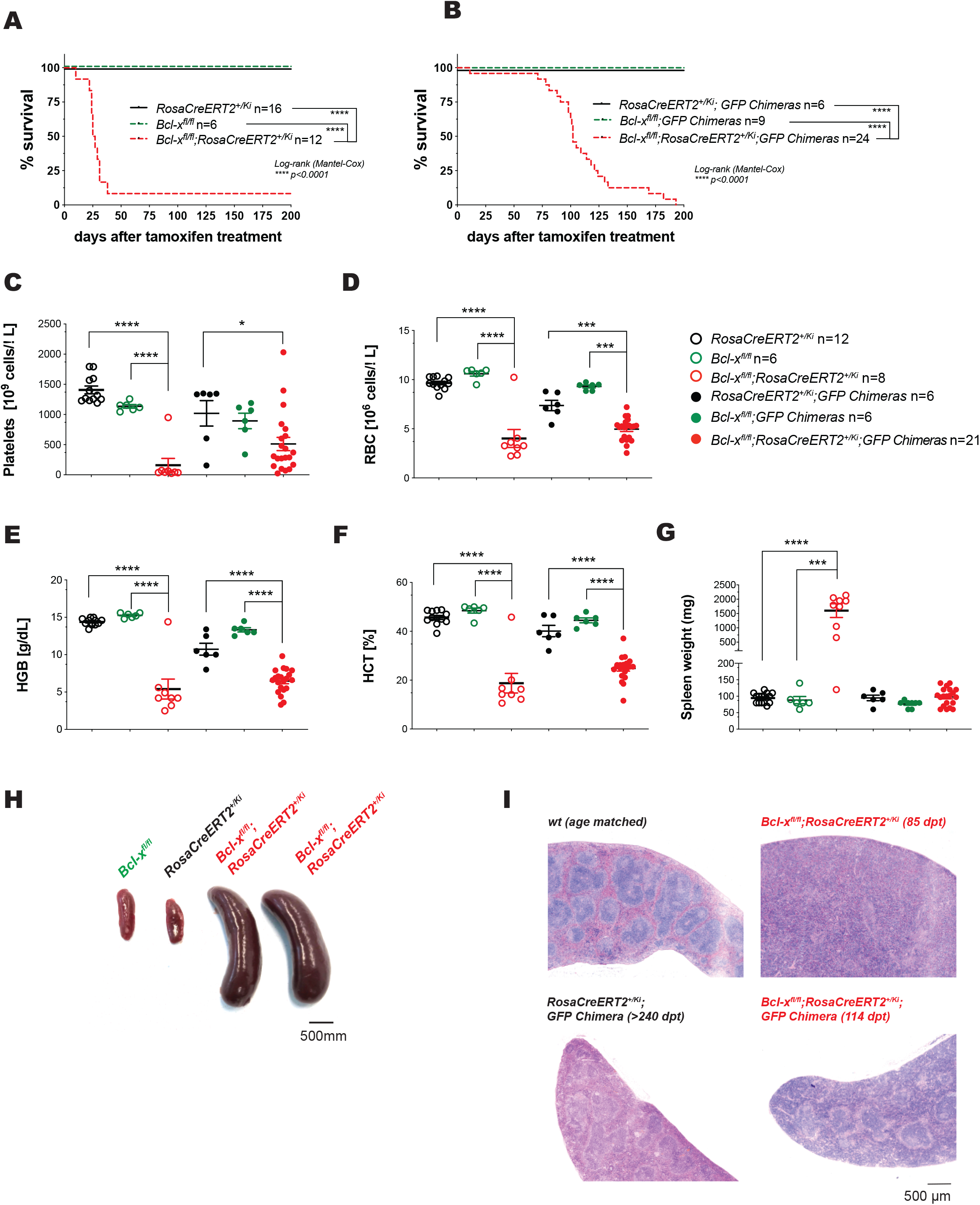
The inducible deletion of BCL-XL causes severe anemia even in chimeric mice that are BCL-XL sufficient in all hematopoietic cell populations. **(A)** *Bcl-x*^*fl*/*fl*^;*RosaCreERT2*^+/*Ki*^ or, as controls, *Bcl-x*^*fl*/*fl*^ *and RosaCreERT2*^+/*Ki*^ mice (age 9-12 weeks, males and females, numbers indicated) were treated with tamoxifen (200 mg/kg/body weight administered in 3 daily doses by oral gavage) to induce CreERT2-mediated deletion of the floxed *Bcl*-*x* alleles. Mice were monitored for up to 200 days post-treatment with tamoxifen. **(B)** *Bcl-x*^*fl*/*fl*^;*RosaCreERT2*^+/*Ki*^ or, as controls, *Bcl-x*^*fl*/*fl*^ *and RosaCreERT2*^+/*Ki*^ mice (males and females, aged 8-14 weeks, numbers indicated in the figure legend) were lethally γ-irradiated (2 × 5.5 Gy, 3 h apart) and reconstituted with bone marrow from UBC-GFP mice (referred to as GFP-Chimeras). After 8 weeks, reconstituted mice were treated with tamoxifen (200 mg/kg body weight administered in 3 daily doses oral gavage) and monitored for up to 200 days (termination of the experiment). **(A-B)** Data are presented as % survival post-treatment with tamoxifen and statistical significance was assessed using the Mantel-Cox (Log-rank) test; ****p<0.0001. **(C)** Total counts of platelets, **(D)** red blood cells (RBC), **(E)** hemoglobin (HGB) content and **(F)** hematocrit (HCT) of tamoxifen-treated mice were determined by ADVIA. **(G)** Spleen weights were measured in sick mice (at sacrifice) or, for the healthy control mice, at the termination of the experiment. **(C-G)** Data are presented as mean ±SEM. Each data point represents an individual mouse and numbers are indicated. Statistical significance was assessed using the Student’s t-test; ****p<0.0001. **(H)** Representative image of enlarged spleens from two *Bcl-x*^*fl*/*fl*^;*RosaCreERT2*^*Ki*/+^ mice and age-matched control *Bcl-x*^*fl*/*fl*^ and *RosaCreERT2*^+/*Ki*^ mice (34 days after treatment). **(I)** Histological analysis of H&E-stained sections of spleens of tamoxifen-treated mice of the indicated genotypes.

To identify the essential functions of BCL-XL in non-hematopoietic tissues of adult mice we generated bone marrow chimeras: *Bcl-x*^*fl*/*fl*^;*RosaCreERT2*^+/*Ki*^ mice that had been lethally irradiated and reconstituted with wild-type bone marrow cells from UBC-GFP mice (GFP-Chimeras; experimental design in Supplementary Figure S1B). Unexpectedly, these animals also presented with severe anemia and thrombocytopenia after loss of BCL-XL, albeit at a considerably later time (Figure 1C-F). Around day 100 post-treatment with tamoxifen, the *Bcl-x*^*fl*/*fl*^;*RosaCreERT2*^+/*Ki*^;*GFP-Chimeras* appeared runty, lethargic and anemic and were sacrificed after >15% body weight loss (median survival = 102 days, Figures 1B). Notably, sick *Bcl-x*^*fl*/*fl*^;*RosaCreERT2*^+/*Ki*^;*GFP-Chimeras* did not present with splenomegaly (Figure 1G) or an increase in the red pulp area of the spleen (Figure 1I), which are hallmarks of primary anemia caused by loss of BCL-XL in erythroid and megakaryocytic cell populations (Figure 1G-I). Importantly, the tamoxifen-treated *Bcl-x*^*fl*/*fl*^;*GFP-Chimera* and *RosaCreERT2*^+/*Ki*^;*GFP-Chimera* control mice all survived without any adverse events for >240 days (termination of the experiment).

These findings demonstrate that inducible loss of BCL-XL in all tissues of adult mice (including hematopoietic cells) causes primary anemia and thrombocytopenia, whereas the combination of total body γ-irradiation (TBI), followed by rescue with a UBC-GFP hematopoietic system, and inducible loss of BCL-XL only in non-hematopoietic cells causes secondary anemia.

### The secondary anemia in mice caused by the combination of γ-radiation and inducible deletion of BCL-XL is not driven by inflammation, hematopoietic malignancy or liver damage

TBI is commonly used for the treatment of hematological malignancies, often alongside with high dose chemotherapy, prior to hematopoietic stem/progenitor cell (HSPC) transplantion. TBI causes many side-effects, including nausea, diarrhoea, sensitive skin and hair loss, that are temporary. However, TBI can also cause severe long-term damage to several organs, such as the lung, trachea and mouth (radiation-induced pneumonitis, fibrosis, oral mucositis) (Marks et al., 2003; Mehta, 2005), reproductive system (radiation-induced infertility) (Ogilvy-Stuart and Shalet, 1993), liver (radiation-induced liver disease) (Kim and Jung, 2017), gastro-intestinal tract (radiation-induced gastric mucositis, radiation-induced gastro intestinal syndrome) (Francois et al., 2013; Olcina and Giaccia, 2016) or kidney (radiation-induced nephropathy) (Cohen, 2000; Cohen and Robbins, 2003). Furthermore, TBI can also initiate secondary malignancies (Dracham et al., 2018).

The above-mentioned TBI-induced pathologies frequently cause secondary anemia (Davis and Littlewood, 2012; Weiss and Goodnough, 2005). For example, hematological malignancies impair the production of RBCs in the bone marrow as a consequence of competing for essential growth factors, the production of reactive oxygen species (ROS) and pro-inflammatory cytokines that damage erythroid progenitors (Davis and Littlewood, 2012; Weiss and Goodnough, 2005). Moreover, a robust inflammatory response is observed upon radiation-induced damage to the gastro-intestinal tract (Olcina and Giaccia, 2016). Pro-inflammatory cytokines (e.g. IL-6, TNFα) interfere with the production of erythropoietin (EPO) and the availability of iron, both crucial for RBC development. Furthermore, inflammatory cytokines enhance the production of white blood cells (WBCs) and can thereby decrease the differentiation of progenitors into RBCs (Weiss and Goodnough, 2005). We did not detect signs of ongoing inflammation as shown by the observation that the numbers of WBCs (Figure 2A), including lymphocytes, neutrophils, basophils, eosinophils and monocytes, were all within the normal range (Supplementary Figure S2A). The neutrophil to lymphocyte ratio (NLR), even though increased in the tamoxifen-treated *Bcl-x*^*fl*/*fl*^;*RosaCreERT2*^+/*Ki*^;*GFP-Chimeras,* was still within the normal range (Figure 2B). Moreover, no signs of hematological malignancies, such as increased numbers of leukocytes were observed in the blood or bone marrow of the tamoxifen-treated *Bcl-x*^*fl*/*fl*^;*RosaCreERT2*^+/*Ki*^;*GFP-Chimeras* (Figure 2A and C).

**Figure 2:**
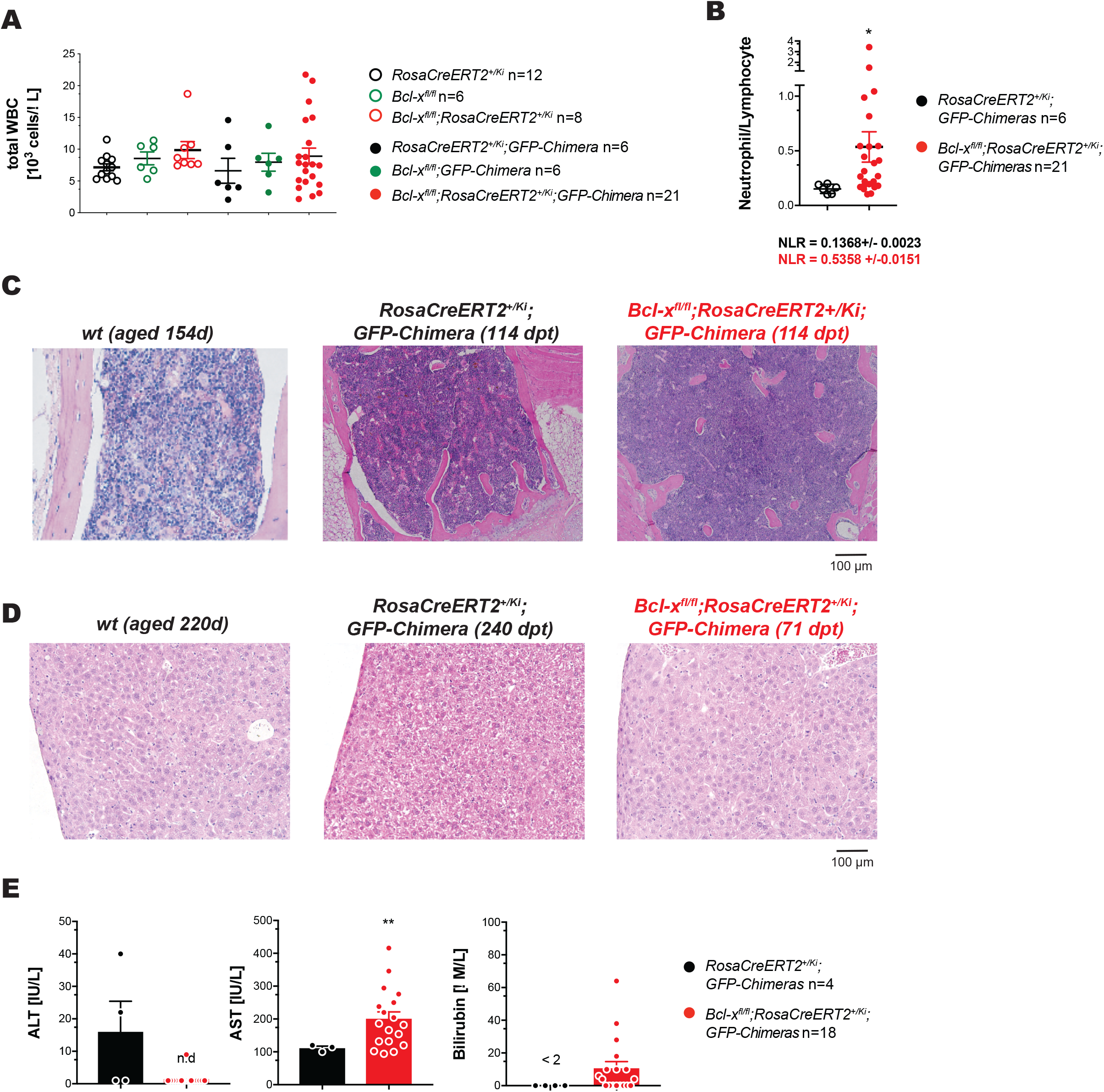
The combination of γ-radiation and inducible deletion of BCL-XL causes neither inflammation, hematopoietic malignancy nor liver damage. *Bcl-x*^*fl*/*fl*^;*RosaCreERT2*^+/*Ki*^ (n=8) or, as controls, *Bcl-x*^*fl*/*fl*^ (n=6) *and RosaCreERT2*^+/*Ki*^ (n=12) mice as well as *Bcl-x*^*fl*/*fl*^;*RosaCreERT2*^+/*Ki*^;*GFP-chimeras* (n=21), or, as controls *Bcl-x*^*fl*/*fl*^;*GFP-chimeras* (n=6) *and RosaCreERT2*^+/*Ki*^;*GFP-chimeras* (n=6) (age 8-14 weeks, males and females) were treated with tamoxifen (200 mg/kg/body weight administered in 3 daily doses by oral gavage) to induce CreERT2-mediated deletion of the floxed *Bcl*-*x* alleles. **(A)** Total white blood cell counts (WBC) were analyzed by ADVIA in sick mice or at the termination of the experiment (healthy control mice). Data are presented as mean ±SEM. Each data point represents an individual mouse and n numbers are indicated. No statistically significant differences were observed. **(B)** The neutrophil/lymphocyte ratio (NLR) is presented as mean ±SEM for sick *Bcl-x*^*fl*/*fl*^;*RosaCreERT2*^+/*Ki*^;*GFP-chimeras* (n=21) or healthy control *RosaCreERT2*^+/*Ki*^;*GFP-chimeras* (n=6) at the termination of the experiment. Data are presented as mean ±SEM. Each data point represents an individual mouse. Statistical significance was assessed using the Student’s t-test; *p<0.05. **(C)** Histological analysis of H&E-stained sections of the sternum of sick *Bcl-x*^*fl*/*fl*^;*RosaCreERT2*^+/*Ki*^;*GFP-chimeras* or healthy wild-type and *RosaCreERT2*^+/*Ki*^;*GFP-chimera* control mice at the indicated time points post-treatment with tamoxifen. (dpt=days post treatment). **(D)** Histological analysis of H&E-stained sections of the livers of sick mice or age-matched healthy control wild-type mice or healthy *RosaCreERT2*^+/*Ki*^;*GFP-chimeras* at the termination of the experiment (dpt=days post-treatment). **(E)** ALT (panel), AST (left middle panel) and bilirubin levels (right panel) in the serum were determined in sick *Bcl-x*^*fl*/*fl*^;*RosaCreERT2*^+/*Ki*^;*GFP-chimeras* (n=18) or in healthy *RosaCreERT2*^+/*Ki*^;*GFP-chimeras* (n=4) at the termination of the experiment. Data are presented as mean ±SEM. Each data point represents one individual mouse. Statistical significance was assessed using Student’s t-test; **p<0.01. nd=not detected.

The liver represents the primary storage site of iron and source of hepcidin, both essential for RBC-production. Hence, secondary anemia can result from liver damage. Histological analysis failed to reveal any abnormalities in the livers of the tamoxifen-treated *Bcl-x*^*fl*/*fl*^;*RosaCreERT2*^+/*Ki*^;*GFP-Chimeras* (Figure 2D) and no marked increases in the liver enzyme ALT or unconjugated bilirubin were detected in their sera (Figure 2E). However, these animals presented with increased levels of AST (Figure 2E), but unlike ALT, AST is also found in the heart, skeletal muscle, kidney and RBCs. Thus, with the lack of other indicators of liver pathology, we attributed this AST-increase to non-liver toxicity.

These findings demonstrate that the secondary anemia in the tamoxifen-treated *Bcl-x*^*fl*/*fl*^;*RosaCreERT2*^+/*Ki*^;*GFP-Chimeras* is not a consequence of erythroid cell destruction, hematologic malignancy, chronic infection, inflammation nor liver damage. Instead, these results suggest that the secondary anemia might be caused by radiation-induced kidney damage.

### The combination of γ-radiation and inducible deletion of BCL-XL causes severe kidney damage in adult mice

EPO stimulates erythropoiesis and is produced in response to cellular hypoxia in the kidney by interstitial fibroblasts adjacent to the renal proximal convoluted tubule and peritubular capillaries. Acute and chronic kidney disease that damage these cells decreases the production of EPO, thereby compromising the generation of erythroid progenitors (CFU-e) and thus RBC-production. We hypothesized that the secondary anemia observed in tamoxifen-treated *Bcl-x*^*fl*/*fl*^;*RosaCreERT2*^+/*Ki*^;*GFP-Chimeras* results from damage to the EPO-secreting cells in the kidney. Accordingly, urine tests showed severe proteinuria in sick tamoxifen-treated *Bcl-x*^*fl*/*fl*^;*RosaCreERT2*^+/*Ki*^;*GFP-Chimeras* (Supplementary Figure S2F) and due to increased urination the bedding of their cages had to be changed frequently.

The kidneys of these sick mice were abnormally small and yellow (Figure 3A and B). Histological analysis revealed severe renal tubulo-interstitial disease with segmental areas showing diminution of the renal tubules, interstitial fibrosis and patchy chronic inflammation with focal and segmental glomerular changes and thickening of renal arterioles. The renal medulla and papillae contained intraluminal calcified cellular debris and crystalline precipitates within the collecting ducts and dilated loops of Henle, where urine is concentrated (Figure 3C). This was sometimes accompanied by intraluminal multinucleated macrophages, reminiscent of the changes seen in myeloma nephropathy. The glomeruli in the affected segments showed secondary focal and segmental sclerosis. These structural changes were additionally visualized using confocal microscopy (Supplementary Figure S3).

**Figure 3:**
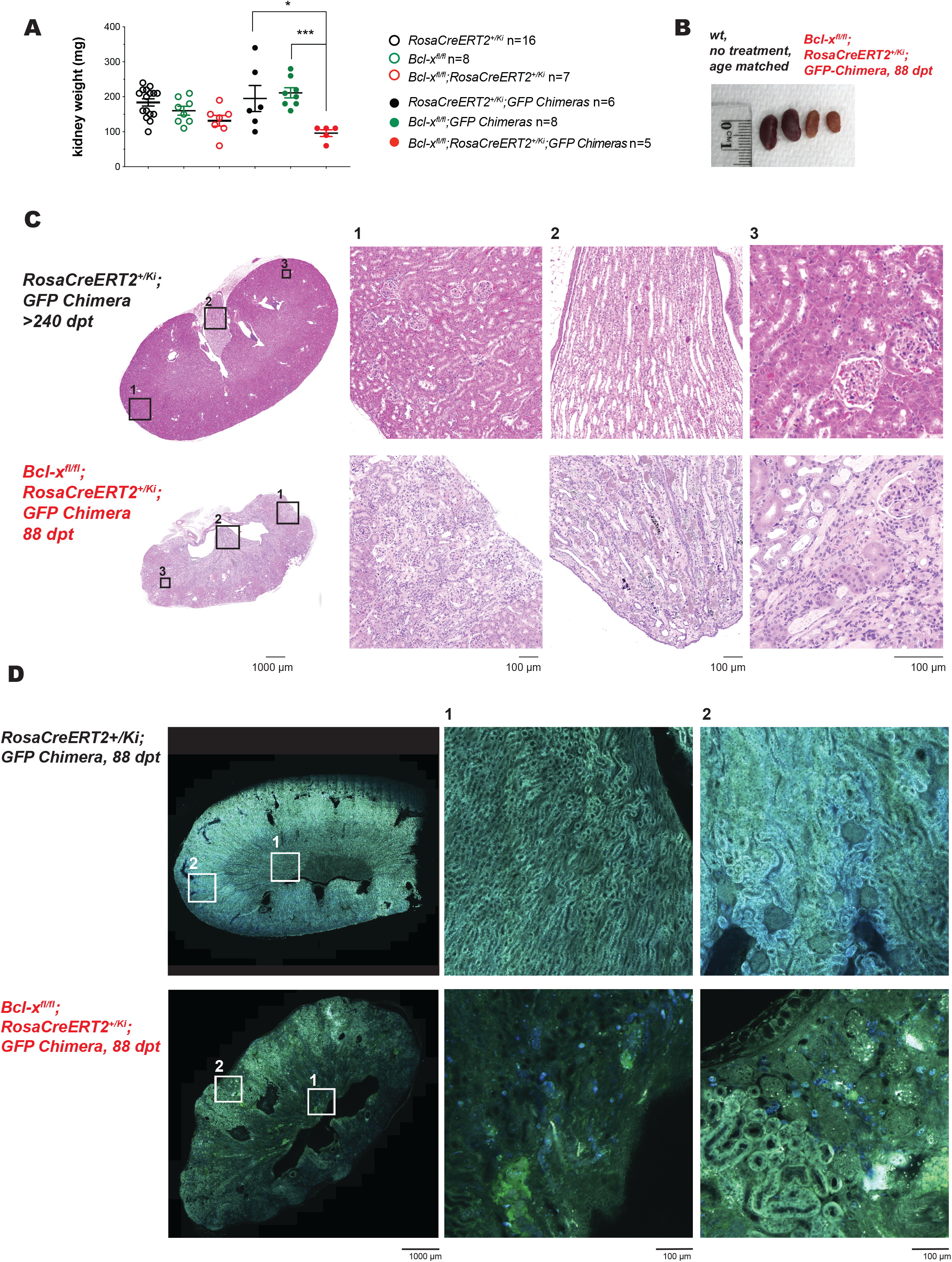
The combination of γ-radiation and inducible deletion of BCL-XL causes severe kidney damage. *Bcl-x*^*fl*/*fl*^;*RosaCreERT2*^+/*K*i^ (n=7) or, as controls, *Bcl-x*^*fl*/*fl*^ (n=8) and *RosaCreERT2*^+/*Ki*^ (n=16) mice as well as *Bcl-x*^*fl*/*fl*^;*RosaCreERT2*^+/*Ki*^;*GFP-chimeras* (n=5), *Bcl-x*^*fl*/*fl*^;*GFP-chimeras* (n=8) and *RosaCreERT2*^+/*Ki*^;*GFP-chimeras* (n=6) (age 8-14 weeks, males and females) were treated with tamoxifen (200 mg/kg/body weight administered in 3 daily doses by oral gavage) to induce CreERT2-mediated deletion of the floxed *Bcl*-*x* alleles. **(A)** Kidney weights were measured in sick *Bcl-x*^*fl*/*fl*^;*RosaCreERT2*^+/*Ki*^;*GFP-chimeras* (at sacrifice) or, for the healthy control mice, at the termination of the experiment. Data are presented as mean ±SEM. Each data point represents one individual mouse. Statistical significance was assessed using Student’s t-test; *p<0.05, ***p<0.001. **(B)** Picture showing both kidneys from age-matched healthy control wild-type mice (left) and sick *Bcl-x*^*fl*/*fl*^;*RosaCreERT2*^+/*Ki*^;*GFP-chimeras* (right). **(C)** Histological analysis of H&E-stained sections of the kidneys of age-matched healthy control mice or sick *Bcl-x*^*fl*/*fl*^;*RosaCreERT2*^+/*Ki*^;*GFP-chimeras* as indicated, showing from left to right, shrunken scarred kidney, segmental chronic tubulo-interstitial disease, accumulation of cellular debris and amorphous material within the collecting ducts of the papilla. The higher magnification shows tubular epithelial degeneration, apoptosis and segmental secondary glomerular sclerosis. Pictures are representative of at least 5 mice for each genotype. **(D)** Multiphoton analysis of fixed sections of the kidney of age-matched healthy *RosaCreERT2*^+/*Ki*^;*GFP-chimeras* (left) or sick *Bcl-x*^*fl*/*fl*^;*RosaCreERT2*^+/*Ki*^;*GFP-chimeras* (right). Pictures are representative of at least 3 mice for each genotype. green=second harmony, blue=short wavelength autofluorescence.

The kidney damage was confirmed using multiphoton microscopy (Figure 3D). To avoid spurious signals due to non-specific antibody binding, the kidneys were visualized using short wavelength auto-fluorescence (blue) and second harmony generation (SHG) (green). Materials that efficiently generate SHG include type-1-collagen, myosin, tubulin or non-linear crystals and minerals. Intensity of short wavelength auto-fluorescence provides information on the metabolic activity as it visualizes metabolic co-factors, including NADH and NADPH. Accordingly, higher short wavelength auto-fluorescence was observed in the mitochondria-rich proximal tubular cells compared to the distal tubular cells in the kidneys from healthy control mice. In contrast, a significant reduction in metabolically active cells was observed in the atrophic segmental areas in sick mice (Figure 3D). The above-mentioned precipitates within the dilated loops of Henle generated a strong SHG signal, confirming their crystalline nature (Figure 3D).

These findings demonstrate that the combination of TBI and *Bcl-x*-deletion cause fatal renal tubulo-interstitial disease.

### The combination of γ-radiation and inducible deletion of BCL-XL causes apoptosis of tubular epithelial cells in the kidney

Loss or inhibition of pro-survival BCL-XL causes apoptotic cell death (Adams and Cory, 2007). We therefore performed TUNEL staining that detects 3′-hydroxyl termini in double-strand DNA breaks, a hallmark of apoptotic cells (Gavrieli et al., 1992; Gorczyca et al., 1992), to determine whether inducible loss of BCL-XL, TBI or their combination caused apoptosis in renal cells. We observed significantly increased numbers of TUNEL^+^ cells in kidney sections from sick tamoxifen-treated *Bcl-x*^*fl*/*fl*^;*RosaCreERT2*^+/*Ki*^;*GFP-Chimeras*; they were mainly localized in the proximal tubular epithelium within segments showing tubulo-interstitial disease. In contrast, only occasional apoptotic cells were found in sections from control animals (Figure 4A and B).

**Figure 4:**
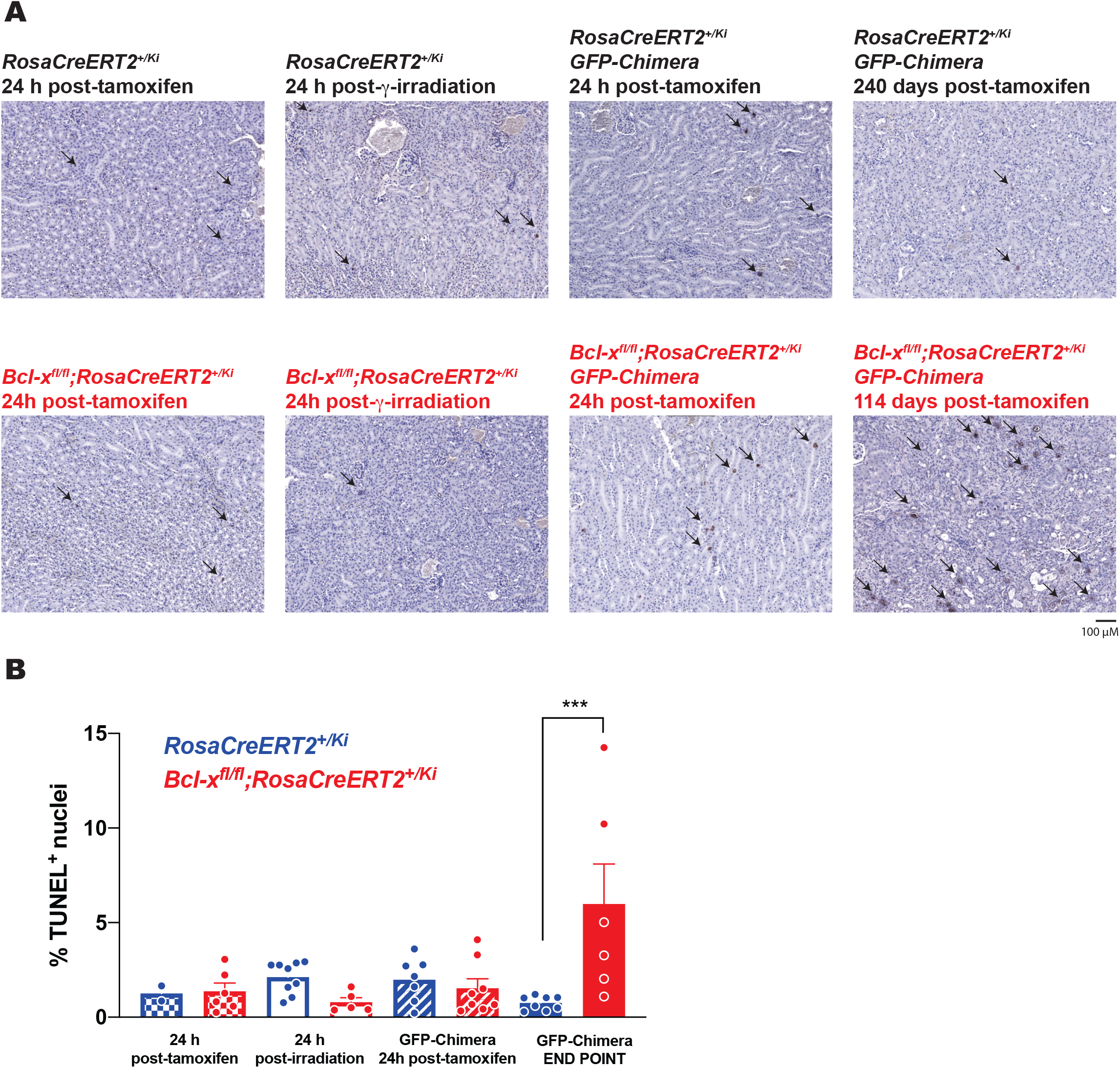
The combination of γ-radiation and inducible deletion of BCL-XL causes apoptosis of proximal renal tubule epithelial cells. **(A)** TUNEL staining of kidney sections from control *RosaCreERT2*^+/*Ki*^, *Bcl-x*^*fl*/*fl*^;*RosaCreERT2*^+/*Ki*^, *RosaCreERT2*^+/*Ki*^;*GFP-chimeras* or *Bcl-x*^*fl*/*fl*^;*RosaCreERT2*^+/*Ki*^;*GFP-chimeras* at the indicated time points post-tamoxifen treatment (200 mg/kg/body weight administered in 3 daily doses by oral gavage) or γ-irradiation (2 × 550 Rad), as indicated. Slides were counterstained with hematoxylin. Arrow heads indicate examples of TUNEL positive (apoptotic) cells. **(B)** TUNEL positive cells were quantified using a personalized script for the Imaga J software. Kidney sections were divided into ~40 microscopic images and the total numbers of blue and brown (TUNEL positive) nuclei were determined. The percentages of TUNEL positive nuclei were calculated for each picture. The average value for each kidney section was then determined and plotted as a single data point. Error bars represent SEM from at least 3 independent samples for each genotype. Statistical significance was assessed using one-way ANOVA analysis with Tukey’s multiple comparisons test (comparing control *RosaCreERT2*^+/*Ki*^ with *Bcl-x*^*fl*/*fl*^;*RosaCreERT2*^+/*Ki*^ and control *RosaCreERT2*^+/*Ki*^;*GFP-chimeras* with *Bcl-x*^*fl*/*fl*^;*RosaCreERT2*^+/*Ki*^;*GFP-chimeras.* ***p<0.001

Notably, 24 h following tamoxifen-treatment (8 weeks post-TBI), TUNEL^+^ cells were detected in *Bcl-x*^*fl*/*fl*^;*RosaCreERT2*^+/*Ki*^;*GFP-Chimeras* as well as in control *RosaCreERT2*^+/*Ki*^;*GFP-Chimeras*, *Bcl-x*^*fl*/*fl*^;*RosaCreERT2*^+/*Ki*^ and *RosaCreERT2*^+/*Ki*^ mice (Figure 4A and B). This indicates that both TBI and loss of BCL-XL by themselves induce renal tubular epithelial cell apoptosis. However, in contrast to the control mice that were able to recover from TBI-induced kidney damage, the additional loss of BCL-XL resulted in more extensive apoptosis in the renal tubular epithelium with their cellular debris accumulating in the renal loops of Henle and collecting ducts. This caused secondary obstruction, glomerulopathy and interstitial nephropathy with damage to the adjacent EPO-secreting peritubular interstitial cells.

These findings demonstrate that the combination of TBI and inducible loss of BCL-XL causes extensive apoptosis in the renal proximal convoluted tubular epithelium, leading to chronic renal failure and secondary anemia.

### Concomitant loss of BIM or PUMA delays kidney disease with secondary anemia caused by radiation and the induced loss of BCL-XL in adult mice

We next investigated which BH3-only protein was critical for apoptosis in the renal tubular epithelium. We focused on BIM and PUMA because both play critical roles in DNA damage-induced apoptosis in diverse cell types (Bouillet et al., 1999a; Erlacher et al., 2005; Villunger et al., 2003). We observed a significant extension of survival of tamoxifen-treated *Bcl-x*^*fl*/*fl*^;*RosaCreERT2*^+/*Ki*^;Bim^+/−^;*GFP-Chimeras* (median survival 159 days; p<0.0001), *Bcl-x*^*fl*/*fl*^;*RosaCreERT2*^+/*Ki*^;*Bim*^−/−^;*GFP-Chimeras* (221 days; p<0.0001), *Bcl-x*^*fl*/*fl*^;*RosaCreERT2*^+/*Ki*^;*Puma*^+/−^;*GFP-Chimeras* (125 days) *Bcl-x*^*fl*/*fl*^;*RosaCreERT2*^+/*Ki*^ ;*Puma*^−/−^;*GFP-Chimeras* (281 days; p<0.0001) compared to the *Bcl-x*^*fl*/*fl*^;*RosaCreERT2*^+/*Ki*^;*GFP-Chimeras* (102 days) (Figure 5A,B). Combined loss of one allele of *Bim* plus one allele of *Puma* (*Bcl-x*^*fl*/*fl*^;*RosaCreERT2*^+/*Ki*^;*Bim*^+/−^;*Puma*^+/−^;*GFP-Chimeras*) resulted in a similar extension of lifespan as seen for mice lacking either both alleles of *Bim* or both alleles of *Puma* (median survival 243 days; p<0.0001) (Figure 5A,B). Nevertheless, all these mice ultimately presented with similar pathology as the tamoxifen-treated *Bcl-x*^*fl*/*fl*^;*RosaCreERT2*^+/*Ki*^;*GFP-Chimeras*, including kidney damage (Figure 5C) and secondary anemia (Figure 5D-F).

**Figure 5:**
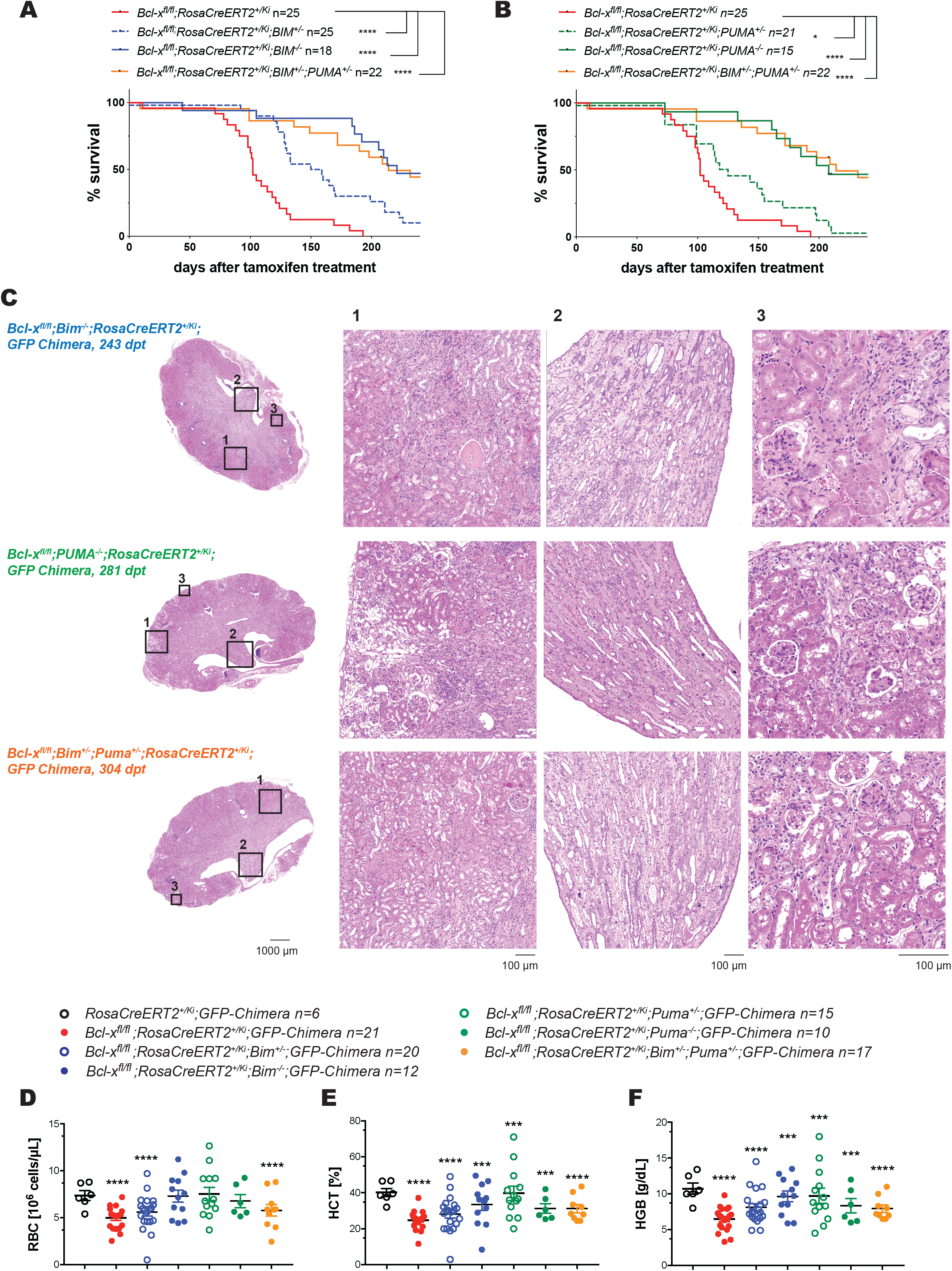
Concomitant loss of BIM or PUMA delays the fatal kidney disease caused by the combination of γ-irradiation and inducible deletion of BCL-XL. Mice of the indicated genotypes (males and females mixed, aged 10-14 weeks, numbers are indicated) were lethally irradiated (2 × 5.5 Gy, 3 h apart) and reconstituted with bone marrow from UBC-GFP mice. After 8 weeks, reconstituted mice were treated with tamoxifen (200 mg/kg body weight administered in 3 daily doses by oral gavage) and monitored for at least 240 days. **(A, B)** Data are presented as % survival and statistical significance was assessed using the Mantel-Cox (Log-rank) test. *p<0.05, ****p<0.0001. The survival curves for the *Bcl-x*^*fl*/*fl*^;*RosaCreERT2*^+/*Ki*^;*GFP-chimeras* are the same as those shown in Figure 1B and presented here for ease of comparison. **(C)** Histological analysis of H&E-stained sections of the kidneys of sick mice of the indicated genotypes at the indicated time points (dpt = days post-tamoxifen treatment). Pictures are representative of at least 3 mice for each genotype. **(D)** Total red blood cell (RBC) count, **(E)** hematocrit (HCT) and **(F)** hemoglobin (HGB) content were determined by ADVIA in the blood of sick mice of the indicated genotypes or at the termination of the experiment for healthy controls. **(D-F)** Data are presented as mean ±SEM. Each data point represents one individual mouse. Numbers of miceare indicated. Statistical significance was assessed using Student’s t-test; ***p<0.001; ****p<0.0001. Data for the *Bcl-x*^*fl*/*fl*^;*RosaCreERT2*^+/*Ki*^;*GFP-chimeras* and *RosaCreERT2*^+/*Ki*^;*GFP-chimeras* are the same as those shown in Figure 1D-F and are shown here for ease of comparison.

These results demonstrate that BIM and PUMA play critical overlapping roles in the induction of apoptosis of renal tubular epithelial cells.

### Pharmacological inhibition of BCL-XL in combination with DNA-damaging anti-cancer therapeutics does not cause kidney disease with secondary anemia

BH3 mimetic drugs targeting select anti-apoptotic BCL-2-family members represent promising novel agents in cancer therapy (Merino et al., 2018). The BCL-XL-specific inhibitor A1331852 (Lessene et al., 2013) has not yet entered clinical trials, mostly due to its on-target toxicity in platelets that is also seen with ABT-263/navitoclax that inhibits BCL-XL, BCL-2 and BCL-W (Leverson et al., 2015; Mason et al., 2007; Wilson et al., 2010). Nevertheless, many cancers depend on BCL-XL for sustained growth (Campbell and Tait, 2018; Merino et al., 2018) and extensive work is being conducted to develop modes of administration and dose schedules for the safe use of BCL-XL inhibitors (Khan et al., 2019). Our present data indicate that the combination of BCL-XL inhibitors with DNA damage-inducing therapies might cause renal toxicity with secondary anemia. However, drug-mediated inhibition of a pro-survival BCL-2 family member often exerts considerably less severe impact than genetic loss of this protein, as demonstrated for MCL-1 (Kotschy et al., 2016). Thus, in contrast to the genetic deletion of *Bcl*-*x* resulting in a permanent loss of BCL-XL protein, the pharmacological inhibition of BCL-XL for a defined period might be tolerated, even in combination with DNA damage-inducing chemotherapeutics. To test this hypothesis, we treated wild-type mice that had been γ-irradiated and reconstituted with UBC-GFP bone marrow with a clinically relevant and well-tolerated dose of the BCL-XL inhibitor A1331852 (Leverson et al., 2015). Additionally, we treated wild-type mice with a combination of clinically relevant and well-tolerated doses of either of the DNA-damaging drugs 5-Fluorouracil (5-FU) or Cyclophosphamide plus A1331852. No adverse effects of any of these treatments were detected for at least 150 days (Figure 6A, B). As expected (Leverson et al., 2015; Mason et al., 2007; Merino et al., 2018; Wilson et al., 2010), treatment with A1331852 resulted in on-target platelet toxicity at day 2 post-treatment, followed by high platelet counts at days 5-6 post treatment (Figure 6C, G). All mice recovered within ~10 days (Figure 6C). This is consistent with the described BCL-XL-dependency of platelets and thus platelet toxicity followed by the rapid induction of a rebound production of platelets upon treatment with BCL-XL-inhibiting BH3 mimetic drugs (Leverson et al., 2015; Mason et al., 2007; Merino et al., 2018; Wilson et al., 2010). As reported (Brinkmann et al., 2017), treatment with 5-FU caused mild anemia (Figure 6E), and mice that were treated with Cyclophosphamide showed a reduction in lymphocytes at day 6 post-treatment (Supplementary Figure S5D). Importantly, for all drug-treated mice, the numbers of RBCs, platelets as well as HGB and HCT values and WBC counts were all in the normal range at the termination of the experiment (Figure 6D and E and Supplementary Figure S4B and D). The *GFP-Chimeras* were slightly lighter and had smaller kidneys and spleens at the end of the experiment (Figure 7A and Supplementary Figure S4A). However, urine testing (at 10 and 100 days post-treatment) did not provide evidence of kidney damage (data not shown). No significant differences in the weights of the kidneys or spleens were observed between mice that were treated with DNA-damaging chemotherapy (5-FU or Cyclophosphamide) alone or in combination with A1331852 when compared to untreated mice (Figure 7B). Histological analysis revealed normal architecture of the kidneys (Figure 7 C,D), liver, spleen and bone marrow in all experimental and control mice (Supplementary Figure S5).

**Figure 6:**
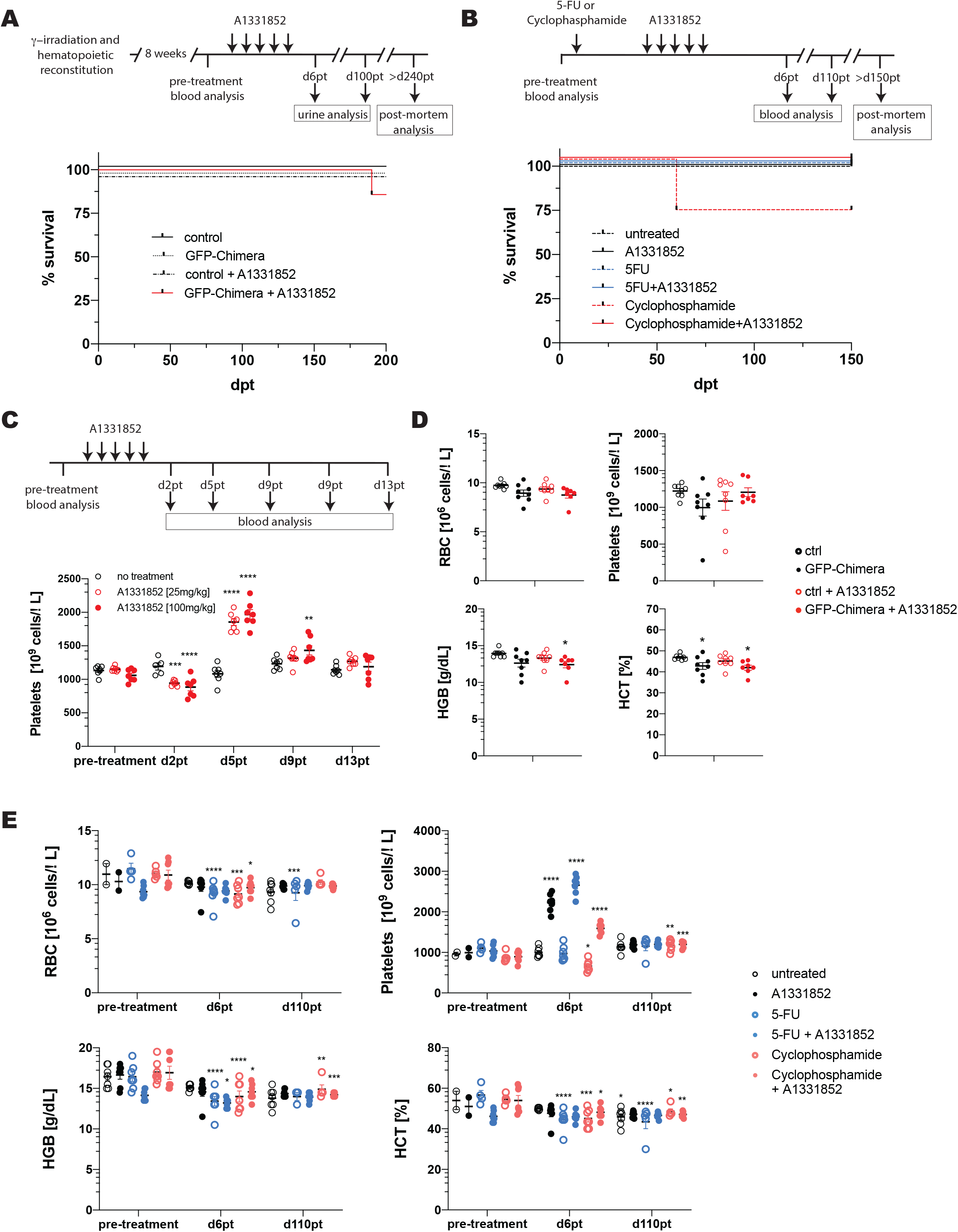
Pharmacological inhibition of BCL-XL in combination with DNA damage-inducing chemotherapeutics or γ-radiation at clinically relevant doses is tolerated in mice. **(A)** GFP-Chimeras were treated with the BCL-XL inhibitor A1331852 (n=8, 5 doses by oral gavage, 25 mg/kg body weight each dose). Control groups (n=8 each group) include untreated GFP-Chimeras and un-irradiated C57BL/6-Ly5.1 (wild-type) mice treated with A1331852 or left untreated. Mice were monitored for up to 200 days post-treatment (dpt). Data are presented as % survival. **(B)** C57BL/6-Ly5.1 (wild-type) mice (females, aged 10 weeks) were treated with Cyclophosphamide (n=7, 150 mg/kg body weight, 1 dose *i.v*.) or 5-Fluorouracil (5-FU, n=7, 100 mg/kg body weight, 1 dose *i.v*.) and after 5 days additionally treated with A1331852 (5 doses by oral gavage, 100 mg/kg body weight each dose). Control groups (n=7 each group) include C57BL/6-Ly5.1 (wild-type) mice (females, aged 10 weeks) treated with A1331852 alone, Cyclophosphamide alone, 5-FU alone or left untreated. Mice were monitored for up to 150 days post-treatment (dpt). Data are presented as % survival. **(C)** C57BL/6-Ly5.1 (wild-type) mice (females, aged 10 weeks) were treated with A1331852 (n=7, 5 doses by oral gavage, 25 or 100 mg/kg body weight each dose). Total platelet counts were determined by ADVIA at the indicated time points. **(D-E)** Total red blood cell count (RBC), platelet count, hemoglobin (HGB) content and hematocrit (HCT) were determined by ADVIA as indicated in the blood of **(D)** drug-treated GFP-Chimeras or control mice (n=8 each group) at the termination of the experiment and **(E)** in drug-treated C57BL/6-Ly5.1 (wild-type) mice or control mice at the indicated time points (n=7 each group). **(C-E)** Data are presented as mean ±SEM. Each data point represents one individual mouse. Statistical significance was assessed using one-way ANOVA analysis with Tukey’s multiple comparisons test (comparing untreated and drug-treated mice within each group). *p<0.05, **p<0.01, ***p<0.001, ****p>0.0001.

**Figure 7:**
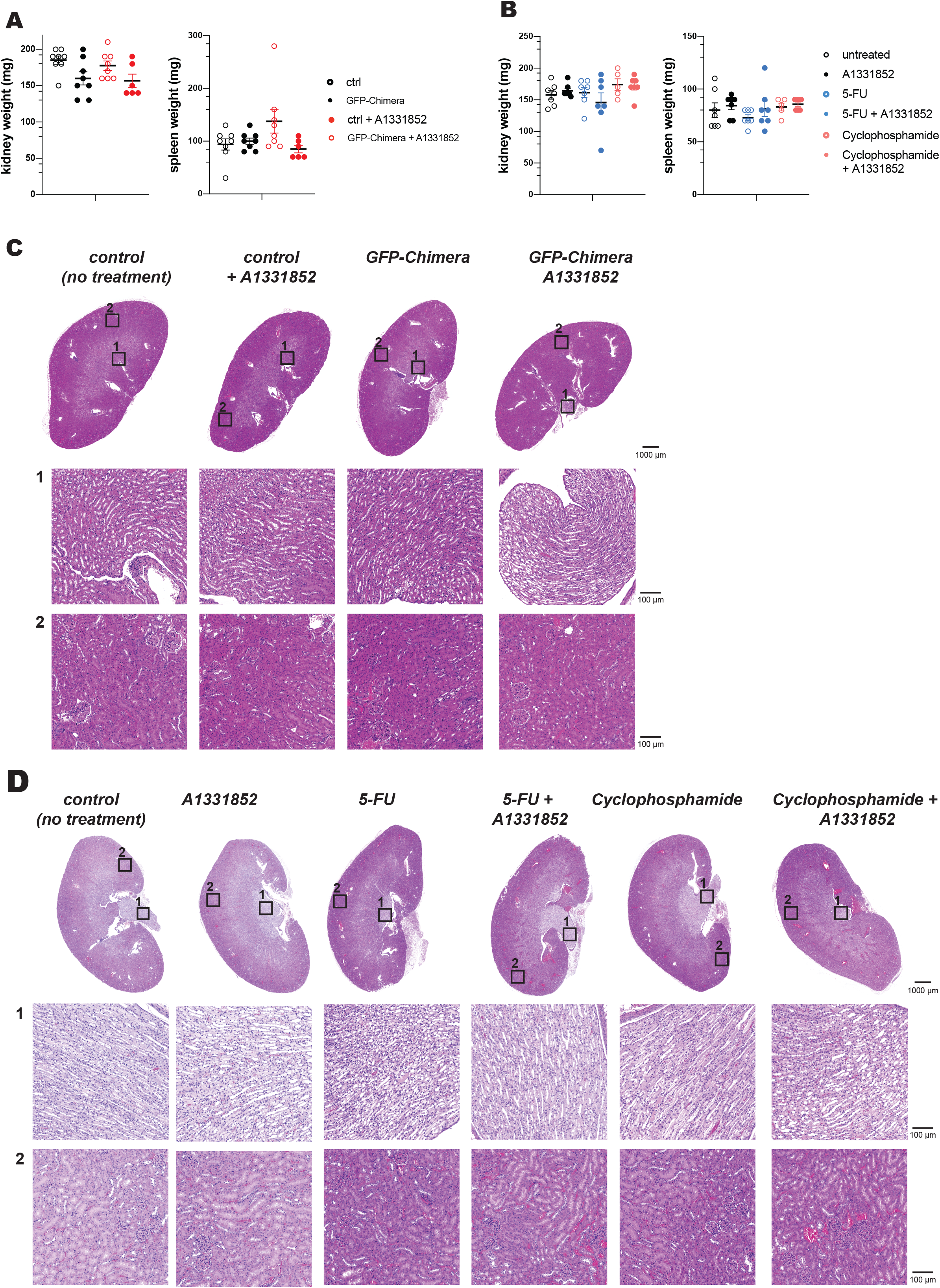
Pharmacological inhibition of BCL-XL in combination with DNA damage-inducing chemotherapeutics or γ-radiation at clinically relevant doses does not impair kidney function and architecture. **(A)** GFP-Chimeras were treated with the BCL-XL inhibitor A1331852 (n=8, 5 doses by oral gavage, 25 mg/kg body weight each dose). Control groups (n=8 each group) include untreated GFP-Chimera and un-irradiated C57BL/6-Ly5.1 (wild-type) mice treated with A1331852 or left untreated. Kidney (left panel) and spleen weights (right panel) were measured in drug-treated GFP-Chimeras or control mice at the termination of the experiment (n=8 each group). **(B)** C57BL/6-Ly5.1 (wild-type) mice (females, aged 10 weeks) were treated with Cyclophosphamide (n=7, 150 mg/kg body weight, 1 dose *i.v*.) or 5-Fluorouracil (5-FU, n=7, 100 mg/kg body weight, 1 dose *i.v*.) and after 5 days additionally treated with A1331852 (5 doses by oral gavage, 100 mg/kg body weight each dose). Control groups (n=7 each group) include C57BL/6-Ly5.1 (wild-type) mice (females, aged 10 weeks) treated with A1331852 alone, Cyclophosphamide alone, 5-FU alone or left untreated. Kidney (left panel) and spleen weights (right panel) were measured in drug-treated mice or control mice at the termination of the experiment (n=7 each group). **(B)** Histological analysis of H&E-stained sections of the kidneys of drug-treated GFP-Chimeras or control mice at the termination of the experiment. Pictures are representative of at least 3 mice for each treatment group. **(C)** Histological analysis of H&E-stained sections of the kidneys of drug-treated mice or control untreated mice at the termination of the experiment. Pictures are representative of at least 3 mice for each treatment group.

These results show that the combination of DNA damage-inducing anti-cancer therapy plus a BCL-XL inhibitor can be tolerated, at least when used sequentially.

## Discussion

TBI and DNA damage-inducing drugs are current mainstays of cancer therapy. Apart from inducing the death of malignant cells, these treatments can also cause damage to diverse healthy tissues, including the kidney, with the underlying mechanisms still not well understood. In mice that received TBI prior to bone marrow transplantion, we found that genetic deletion of BCL-XL exclusively in non-hematopoietic cells resulted in secondary anemia due to kidney damage. This could be substantially delayed by the concomitant loss of pro-apoptotic PUMA or BIM. This reveals that in the context of DNA damage, BCL-XL is critical for the sustained survival of renal tubular epithelial cells in the adult. Notably, BCL-2 loss results in severe polycystic kidney disease that starts during embryonic development (in the absence of DNA damage)(Bouillet et al., 2001; Veis et al., 1993). This indicates that BCL-2 and BCL-XL are both required to maintain renal tubule epithelial cell survival, the former during development and the latter in adulthood.

Radiation-induced nephropathy (RN) is frequently observed in cancer patients receiving TBI and bone marrow transplants (10-25%)(Cohen, 2000; Cohen et al., 2010) (Cohen and Robbins, 2003). Interestingly, only some TBI-patients develop renal injury/failure, and the reasons for this are currently unknown (Cohen and Robbins, 2003). Our findings suggest that BCL-XL is essential to ensure the survival of renal cells that had previously sustained DNA damage, whereas undamaged cells can tolerate the loss of BCL-XL. The reasons for this could be that undamaged cells might express higher levels of pro-survival BCL-2 family members other than BCL-XL (e.g. BCL-2, MCL-1) or that damaged cells express higher levels of pro-apoptotic BH3-only proteins (Campbell and Tait, 2018; Merino et al., 2018). This hypothesis is supported by our demonstration that the concomitant loss of the BH3-only proteins BIM or PUMA can markedly delay kidney damage and secondary anemia caused by the combination of TBI and inducible loss of BCL-XL. Perhaps patients susceptible to RN express higher basal levels of BH3-only proteins in renal tissues, possibly because they are subject to certain stresses already. Alternatively, these cells may express abnormally low levels of pro-survival BCL-2 family members, possibly because they receive lower levels of growth factors (e.g. EGF, FGF).

All structures of the kidney can be affected by RN, including blood vessels, glomeruli, tubular epithelia and the interstitium (Cohen and Robbins, 2003). Our histological analysis indicates that the primary pathology is apoptosis of the proximal renal tubular epithelium rather than endothelial or glomerular damage. Proximal renal tubular epithelium cells are particularly rich in mitochondria and subject to stress due to their constant H^+^-pumping. This might render these cells prone to the induction of apoptosis driven by DNA-damage and the loss of BCL-XL. The deposits within the renal papillae are likely due to the accumulation of apoptotic debris and precipitates from concentrated urine in the loops of Henle or collecting ducts. These secondary obstructive changes are likely to have caused the chronic renal failure (consistent with the observed polyuria) and the segmental cortical scarring and glomerular changes (consistent with proteinuria). The glomerular changes are most likely a secondary effect and not directly caused by TBI as they were not seen in the control *RosaCreERT2*^+/*Ki*^;*GFP-Chimeras*.

Besides TBI, genotoxic (e.g. cisplatin, cytarabine) as well as non-genotoxic drugs (e.g. cyclosporin, methotrexate) can induce kidney damage. With the advent of BH3-mimetic drugs for cancer therapy (Merino et al., 2018), it is important to understand what toxicities might arise from their combination with TBI or standard chemotherapeutics. We found that combinations of the BCL-XL-specific BH3-mimetic A1331852 with TBI, 5-FU or cyclophosphamide were tolerable in mice, at least when used sequentially. This reveals that genetic irreversible loss of *Bcl*-*x* has more severe consequences than pharmacological inhibition of BCL-XL for a defined period. This conclusion is reminiscent of the findings from investigations comparing the impact of genetic loss of pro-survival MCL-1 *vs* treatment of mice for a defined period with an MCL-1 specific BH3-mimetic drug (Caenepeel et al., 2018; Kotschy et al., 2016). These and other studies (Brennan et al., 2018; Brinkmann et al., 2017) suggest that it may be possible to establish therapeutic windows for combination treatment using BH3-mimetic drugs and standard chemotherapy or radiation.

## Materials and Methods

### Mice

*Bclx*^*fl*^ (Wagner et al., 2000), *Bim*^−/−^ (Bouillet et al., 1999b), *Puma*^−/−^ (Villunger et al., 2003), UBC-GFP Tg (Schaefer et al., 2001) and *RosaCreERT2* (Seibler et al., 2003) mice have been described. All mice were either generated on a C57BL/6 background or had been backcrossed onto this background for at least 20 generations. All experiments with mice were conducted according to the guidelines of The Walter and Eliza Hall Institute of Medical Research Animal Ethics Committee.

### Generation of bone marrow chimeras

Recipient mice were subjected to 2 doses of 5.5 Gy given 3 h apart. After 2 h, mice were transplanted with 3-6×10^6^ UBC-GFP Tg bone marrow cells. Successful hematopoietic reconstitution was verified by detecting GFP+ blood cells by FACS.

### Chemotherapeutic drug treatment

To activate CreERT2, mice were given 60 mg/kg tamoxifen (Sigma-Aldrich) in peanut oil/10% ethanol for 3 days by oral gavage 8 weeks post-transplantation. The BCL-XL inhibitor A1331852 was synthesized in house, dissolved in 60% Phosal PG 50, 27.5% polyethylene glycol 400 (PEG-400), 10% ethanol, 2.5% DMSO and administered by oral gavage. Working solutions of 5-Fluorouracil and Cyclophosphamide (Sigma- Aldrich) were prepared according to the manufacturer’s instructions and a maximum volume of 200 μL was injected into the tail vein (*i.v.)*.

### Serum analysis

Sera were stored at −20^°^C and analyzed using Architect c1600 (Abbott Diagnostics,).

### Histology

Tissues were harvested and fixed in 10% formalin. Sections (75 µm) were stained with hematoxylin and eosin (H&E) and examined blinded to the genotype and treatment.

### TUNEL staining

TUNEL staining was performed as described (Salvamoser et al., 2019). Slides were scored by microscopy in a blinded manner. Kidney sections were divided in ~40 microscopic images and the percentage of blue (TUNEL^+^) *vs* brown nuclei was determined using a personalized script for ImageJ. Data points represent average values for each kidney section.

### Microscopy

For multiphoton microscopy kidneys were fixed in 4% paraformaldehyde overnight at 4 °C and embedded in 3% low-melting point agarose (Sigma- Aldrich). Sections (500 µM) were mounted on slides in glycerol. Images were acquired on an Olympus FVMPE-RS Multiphoton system (25x magnification, FV315/5W, NA 1.05, H2O, RT) equipped with a Mai-Tai eHP DeepSee multi-photon laser and processed using CellSens software (Olympus).

For confocal imaging, kidneys were fixed in 10% formalin. Sections (75 µm) were mounted on adherent microscopic slides (Sarstedt). Slides were blocked in 2% normal donkey serum, 1% TritonX100 in PBS for 20 min at RT and stained overnight with rat-anti-PECAM1/CD31 antibodies (1/100, clone Mec13.3, BD Pharmingen) in blocking solution and secondary Alexa594 goat anti-rat IgG antibodies and DAPI (Invitrogen) in blocking solution for 1 h. Slides were coverslipped with ProLong mounting media (Molecular Probes, Invitrogen) and imaged using a Zeiss LSM780 (25x magnicfication, oil, NA 0.8, RT).

### Statistical analysis

Data were plotted and analyzed with Prism (GraphPad Software Inc). Statistical comparisons were conducted using unpaired two-tailed Student’s t test assuming equal variance or one-way ANOVA analysis with Tukey’s multiple comparisons test as indicated. N-numbers for individual experiments are provided in the figure legends.

## Author contributions

KB, SGr, SGl and AS designed the study. MJH helped with the design of experiments. KB conducted all experiments. PW performed pathological analysis of tissue specimens; AD and GK helped with some experiments. VW helped with the multi-photon microscopic analysis. LW performed the quantification of the TUNEL assays. KB, PW and AS wrote the manuscript and all other authors edited it.

## Acknowledgments

We thank Drs P Bouillet, JM Adams, L Hennighausen, and A Villunger for gifts of mice; T. Kitson, C D’Alessandro, C Gatt, S O’Connor, J Mansheim, K McKenzie and G Siciliano for expert animal care; B Helbert for genotyping; J Corbin and J McManus for automated blood analysis. We thank L O Reilly for helping with the TUNEL assays. We thank KA Sarosiek for the helpful discussion of some results. This work was supported by grants and fellowships from the Deutsche Krebshilfe (Dr Mildred Scheel post-doctoral fellowship to KB), Cancer Council of Victoria (SG, AD ‘Sydney Parker Smith Postdoctoral Fellowship’), Leukaemia Foundation Australia (SG), the Lady Tata Memorial Trust (SG), Cure Brain Cancer Australia (AS), the National Health and Medical Research Council (Program Grants #1016701 and 1016647, NHMRC Australia Fellowship 1020363; all to AS), (Project Grants 1145728 to MJH, 1143105 to MJH and AS, and Fellowships 1156095 to MJH) the Leukemia and Lymphoma Society (SCOR Grant #7001-03 to AS), Melbourne International Research and the Melbourne International Fee Remission Scholarship (University of Melbourne, SG) and Cancer Therapeutics CRC Top-up Scholarship (SG, AD). The estate of Anthony (Toni) Redstone OAM, University of Melbourne International Research and International Fee Remission Scholarships (SG), Australian Postgraduate Award (ARDD, JPB), and the operational infrastructure grants through the Australian Government IRIISS and the Victorian State Government OIS.

## Disclosure of Conflict of Interest

The authors declare no conflict of interest.

## Abbreviations

AST: Aspartate aminostransferase
ALT: alanine aminostransferase
CFU-e: colony forming unit-erythroid
EPO: erythropoietin
GIS: gastro intestinal syndrome
HCT: hematocrit
HGB: hemoglobin
IR: γ-irradiation
LDH: lactate dehydrogenase
MOMP: mitochondrial outer membrane permeabilization
NLR: neutrophil to lymphocyte ratio
RBCs: red blood cells
RISM: radiation-induced secondary malignancies
RN: Radiation-induced nephropathy
ROS: reactive oxygen species
SHG: second harmony generation
TBI: total body irradiation
WBCs: white blood cells
5-FU: 5-Fluorouracil

## Notes

### Competing Interest Statement

The authors have declared no competing interest.

