## Supplemental Information for "BCL-XL is essential for the protection from secondary anemia caused by radiation-induced fatal kidney damage"

**Figure S1:** The combination of γ-radiation and inducible deletion of BCL-XL causes fatal secondary anemia in mice

**A**

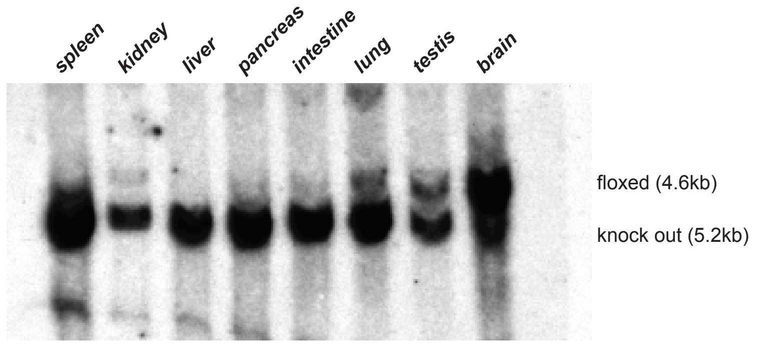


**B**

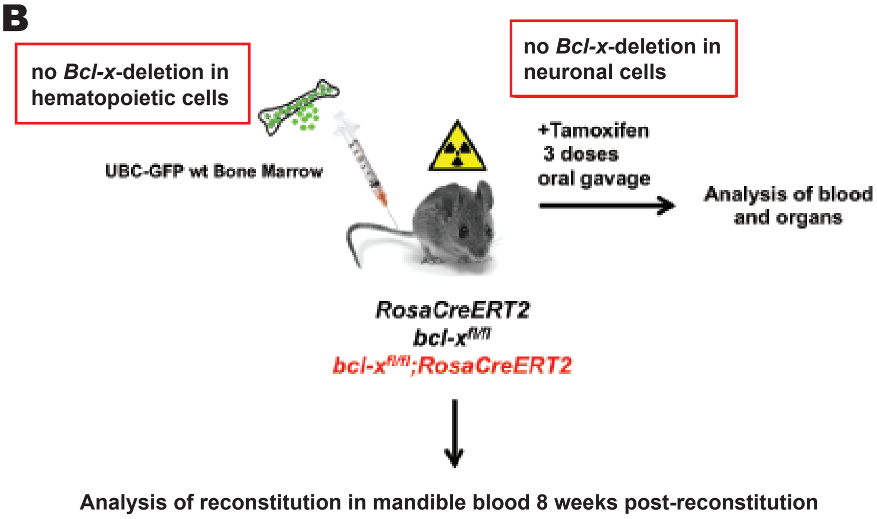


**C**

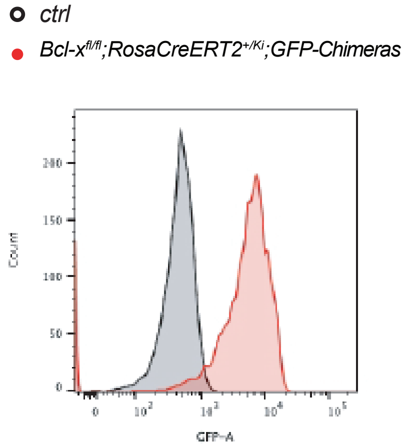

**Figure S2:** The combination of γ-radiation and inducible deletion of BCL-XL neither causes inflammation nor hematopoietic malignancy.

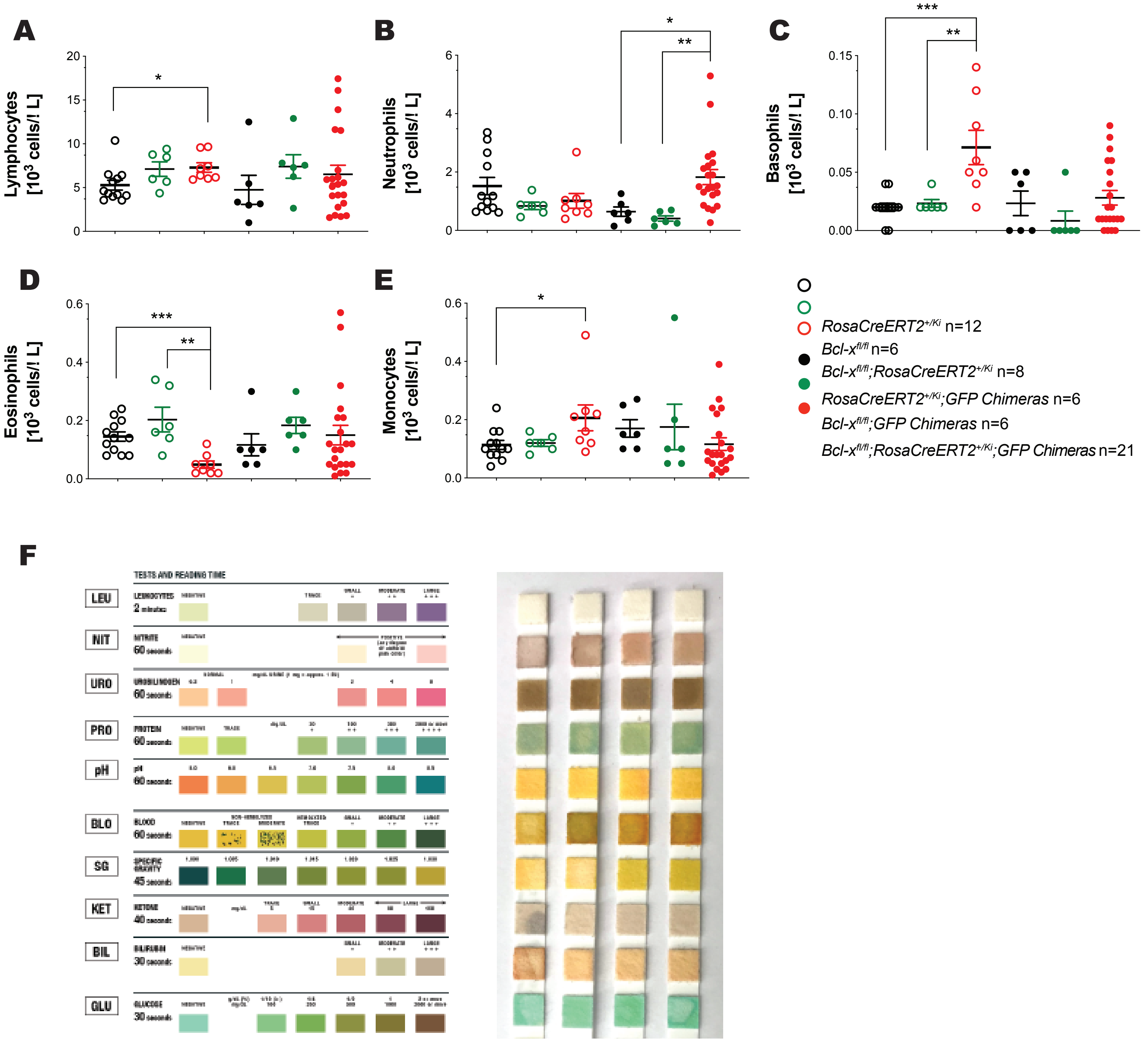

**Figure S3:** The combination of γ-radiation and inducible deletion of BCL-XL causes substantial apoptosis of renal cells and loss of renal architecture.

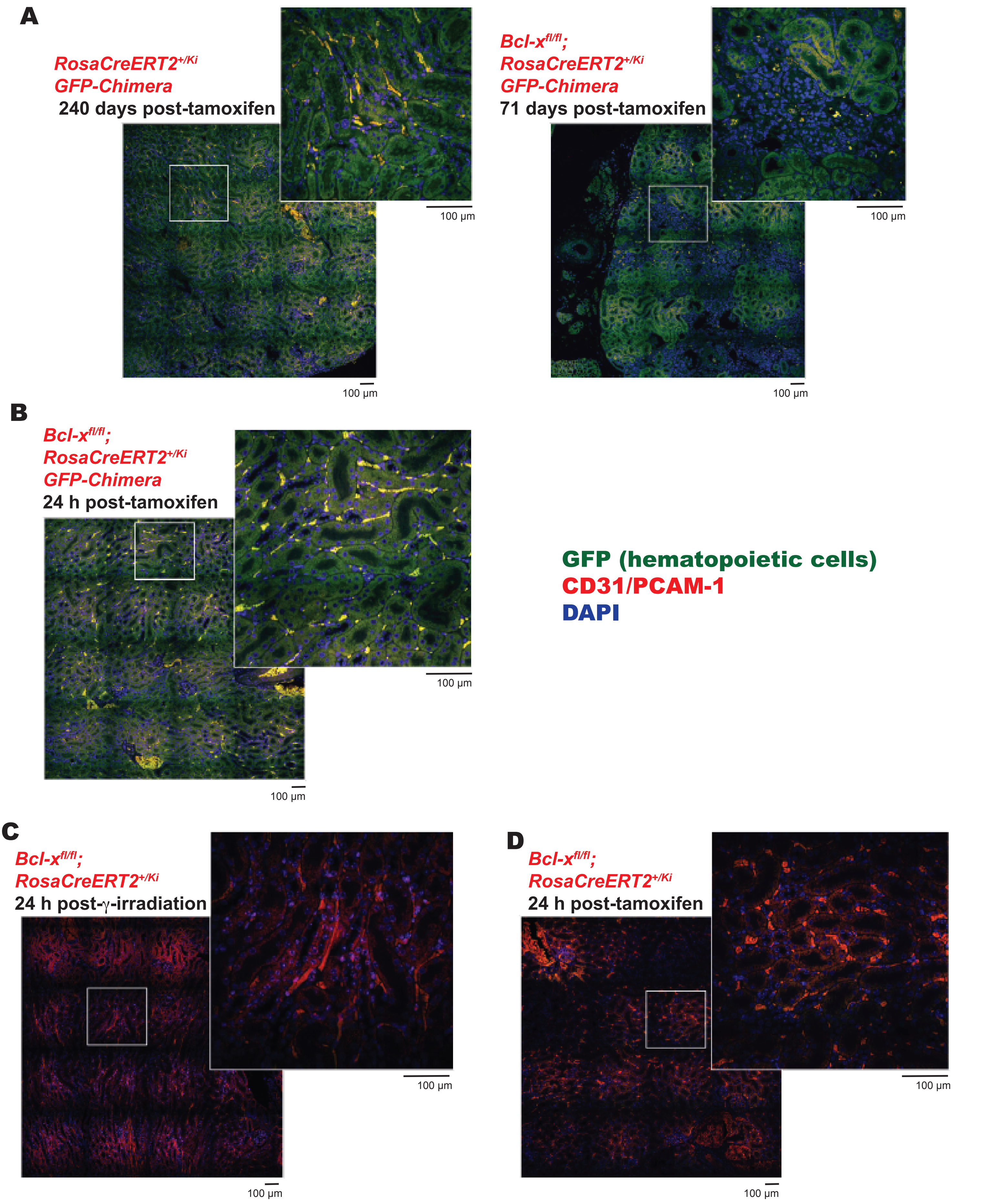

**Figure S4:** Pharmacological inhibition of BCL-XL in combination with DNA damage at clinically relevant doses is tolerated in mice

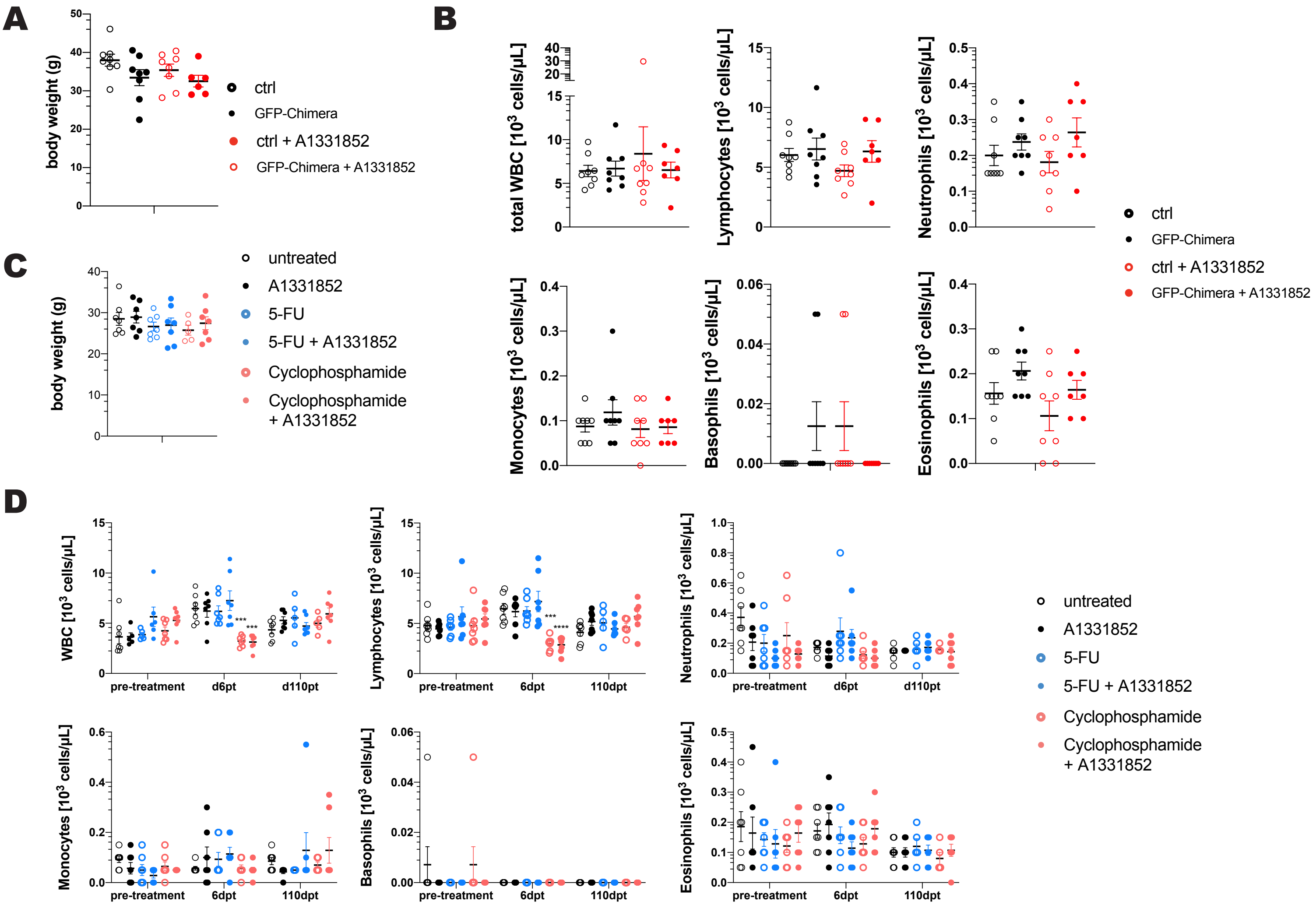

**Figure S5:** Pharmacological inhibition of BCL-XL in combination with DNA damage at clinically relevant doses not impact on normal organ architecture

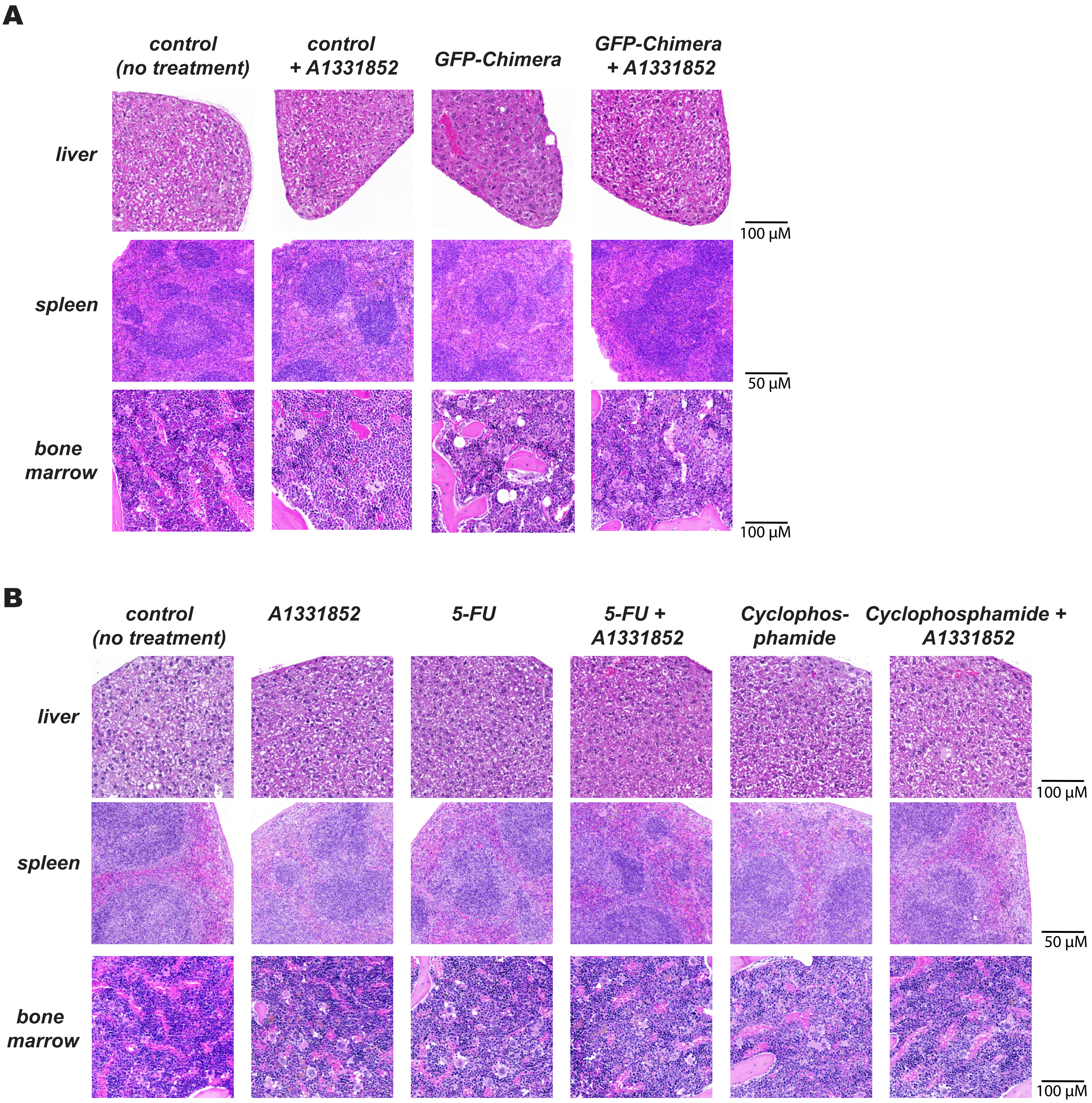

**Supplementary Figure Legends**

**Figure S1: The combination of γ-radiation and inducible deletion of BCL-XL causes secondary anemia in mice. (A)** Southern blot analysis of DNA extracted from the indicated tissues from *Bcl-x$^{fl/fl}$;RosaCreERT2$^{+/Ki}$* mice 24 h after treatment with tamoxifen (200 mg/mouse administered in 5 daily doses by oral gavage). **(B)** Schematic representation of the experimental design. *Bcl-x$^{fl/fl}$;RosaCreERT2$^{+/Ki}$* or, as controls, *Bcl-x$^{fl/fl}$ and RosaCreERT2$^{+/Ki}$* mice (males and females mixed, aged 8-14 weeks) were lethally irradiated (2 x 5.5 Gy, 3 h apart) and reconstituted with bone marrow from UBC-GFP mice (referred to as GFP-Chimeras). After 8 weeks, the reconstitution efficacy was analyzed by measuring GFP+ leukocytes in peripheral blood cells by flow cytometric analysis. Once effective hematopoietic reconstitution was confirmed, the chimeric mice were treated with tamoxifen (200 mg/kg body weight administered in 3 daily doses by oral gavage) and monitored for at least 240 days. **(C)** Flow cytometry histogram showing GFP-MFI (median fluorescence intensity) of total blood derived from non-reconstituted control mice and *Bcl-x$^{fl/fl}$;RosaCreERT2$^{+/Ki}$;GFP-chimeras.*

**Figure S2: The combination of γ-radiation and inducible deletion of BCL-XL causes neither inflammation nor hematopoietic malignancy.** *Bcl-x$^{fl/fl}$;RosaCreERT2$^{+/Ki}$* or, as controls, *Bcl-x$^{fl/fl}$ and RosaCreERT2$^{+/Ki}$* mice (age 9-12 weeks, males and females, n numbers are indicated) were treated with tamoxifen (200 mg/kg/body weight administered in 3 daily doses by oral gavage) to induce CreERT2-mediated deletion of the floxed *Bcl-x* alleles. Mice were monitored for up to 200 days post-treatment with tamoxifen. Another group of *Bcl-x*$^{fl/fl}$*;RosaCreERT2*$^{+/Ki}$ or, as controls, *Bcl-x*$^{fl/fl}$ and *RosaCreERT2*$^{+/Ki}$ mice (males and females, aged 8-14 weeks, numbers indicated in the figure legend) were lethally irradiated (2 x 5.5 Gy, 3 h apart) and reconstituted with bone marrow from UBC-GFP mice (referred to as GFP-Chimeras). After 8 weeks, successfully reconstituted mice were treated with tamoxifen (200 mg/kg body weight administered in 3 daily doses oral gavage) and monitored for up to 200 days. Total numbers of **(A)** lymphoycytes, **(B)** neutrophils, **(C)** basophil, **(D)** eosinophil and **(E)** monocytes were determined by ADVIA in blood from sick mice or at the termination of the experiment for healthy control mice. **(A-E)** Data are presented as mean ±SEM. Each data point represents an individual mouse and numbers of mice are indicated. Statistical significance was assessed using the Student's t-test; $*p<0.05$, $**p<0.01$, $***p<0.001$. **(F)** Representative images of urine strips testing for the indicated markers (Siemens 10SG) in sick *Bcl-x*$^{fl/fl}$*;RosaCreERT2*$^{+/Ki}$*;GFP-Chimeras* (n=4).

**Figure S3: The combination of γ-radiation and inducible deletion of BCL-XL causes substantial apoptosis of renal cells and disturbance of renal architecture.** Confocal microscopic images of kidney sections from **(A)** healthy *RosaCreERT2*$^{+/Ki}$*;GFP-chimeras* at the termination of the experiment (left) or sick *Bcl-x*$^{fl/fl}$*;RosaCreERT2*$^{+/Ki}$*;GFP-chimeras* (right) (dpt=days post-tamoxifen treatment), **(B)** *Bcl-x*$^{fl/fl}$*;RosaCreERT2*$^{+/Ki}$*;GFP-chimeras* 24 h post-tamoxifen treatment, **(C)** *Bcl-x*$^{fl/fl}$*;RosaCreERT2*$^{+/Ki}$ mice 24 h post γ–irradiation (lower panel, left) or **(D)** *Bcl-x*$^{fl/fl}$*;RosaCreERT2*$^{+/Ki}$ mice 24 h post-tamoxifen treatment. Red=CD31/PCAM-1, green=GFP (reconstituted hematopoietic cells), bue=DAPI.

**Figure S4: Pharmacological inhibition of BCL-XL in combination with DNA damage-inducing chemotherapeutics or γ-radiation at clinically relevant doses is tolerated in mice. (A)** GFP-Chimeras were treated with the BCL-XL inhibitor A1331852 (n=8, 5 doses by oral gavage, 25 mg/kg body weight each dose). Control groups (n=8 each group) include untreated GFP-Chimeras and non-irradiated C57BL/6-Ly5.1 (wild-type) mice that had been treated with A1331852 or left untreated. Body weight of drug-treated GFP-Chimeras or intact (non-chimeric) mice was assessed at the termination of the experiment **(B)** Total counts of white blood cells (WBC), lymphocytes, neutrophils, monocytes, basophils and eosinophils were determined by ADVIA as indicated in the blood of drug-treated GFP-Chimeras or control mice at the termination of the experiment. (n=8 each group). **(C)** C57BL/6-Ly5.1 (wild-type) mice (females, aged 10 weeks) were treated with Cyclophosphamide (n=7, 150 mg/kg body weight, 1 dose *i.v.*) or 5-Fluorouracil (5-FU, n=7, 100 mg/kg body weight, 1 dose *i.v.*) and after 5 days additionally treated with A1331852 (5 doses by oral gavage, 100 mg/kg body weight each dose). Control groups (n=7 each group) include C57BL/6-Ly5.1 (wild-type) mice (females, aged 10 weeks) treated with A1331852 alone, Cyclophosphamide alone, 5-FU alone or left untreated. Body weight of drug-treated mice or control mice was assessed at the termination of the experiment **(D)** Total counts of white blood cells (WBC), lymphocytes, neutrophils, monocytes, basophils and eosinophils were determined by ADVIA in the blood of drug-treated mice or control mice at the indicated time points (n=7 each group). **(A-D)** Data are presented as mean ±SEM. Each data point represents one individual mouse. Statistical significance was assessed using Student's t-test; *$p<0.05$.

**Figure S5: Pharmacological inhibition of BCL-XL in combination with DNA damage-inducing chemotherapeutics or γ-radiation at clinically relevant doses does not impact on normal organ architecture (A)** Histological analysis of H&E-stained sections of the liver (upper panel), spleen (middle panel) and bone marrow (sternum) (lower panel) of drug-treated GFP-Chimeras or control mice at the termination of the experiment. Pictures are representative of at least 3 mice for each treatment group. **(B)** Histological analysis of H&E-stained sections of the liver (upper panel), spleen (middle panel) and bone marrow (sternum) (lower panel) of drug-treated mice or healthy control mice at the termination of the experiment, as indicated. Pictures are representative of at least 3 mice for each treatment group.
